# Divergent strategies for learning in males and females

**DOI:** 10.1101/852830

**Authors:** Cathy S. Chen, R. Becket Ebitz, Sylvia R. Bindas, A. David Redish, Benjamin Y. Hayden, Nicola M. Grissom

## Abstract

A frequent assumption in value-based decision-making tasks is that agents make decisions based on the feature dimension that reward probabilities vary on. However, in complex, multidimensional environments, stimuli can vary on multiple dimensions at once, meaning that the feature deserving the most credit for outcomes is not always obvious. As a result, individuals may vary in the strategies used to sample stimuli across dimensions, and these strategies may have an unrecognized influence on decision-making. Sex is a proxy for multiple genetic and endocrine influences that can influence decision-making strategies, including how environments are sampled. In this study, we examined the strategies adopted by female and male mice as they learned the value of stimuli that varied in both image and location in a visually-cued two-armed bandit, allowing two possible dimensions to learn about. Female mice acquired the correct image-value associations more quickly than male mice, and they used a fundamentally different strategy to do so. Female mice constrained their decision-space early in learning by preferentially sampling one location over which images varied. Conversely, male strategies were inconsistent, changing frequently and strongly influenced by the immediate experience of stochastic rewards. Individual strategies were related to sex-gated changes in neuronal activation in early learning. Together, we find that in mice, sex is linked with divergent strategies for sampling and learning about the world, revealing substantial unrecognized variability in the approaches implemented during value-based decision-making.

## Introduction

Value-based decision-making tasks are used by researchers across species to determine the cognitive and neural mechanisms for reinforcement learning and choice behavior [1–4]. One frequent assumption in value-based decision-making tasks, such as bandit tasks, is that the agents make their decisions based on the feature dimension that the experimenter has designed the reward probabilities to vary on, because the other feature dimensions are uninformative and counterbalanced. However, in complex, multidimensional environments, the feature contributing most to reward outcomes is not always obvious to the individual agent making the choices, and stimuli can simultaneously vary on multiple feature dimensions, such as identity and location, at once [5]. As a result of this complexity, differences in learning and decision making within and between individuals over time could result as much from differences in the strategies employed to sample across the different feature dimensions of stimuli, as they could from more straightforward reinforcement learning processes. Because value-based decision tasks are increasingly used to assess and interpret cognition not only in typical situations, but in conditions relevant to neuropsychiatric disease [6–9], recognizing the diverse strategies that can be employed during value based decision making in typical individuals, and the factors that influence strategy selection, is essential.

Rodents, particularly mice, are increasingly employed to elucidate the fundamental neural mechanisms of value-based decision-making [3,10–14]. One under-recognized strength of mouse models is the ease of running many mice simultaneously and repeatedly, allowing the analysis of individual differences between animals in decision strategies. One potential source of phenotypic variance in value-based decision-making that mice are well-suited to studying is the influence of sex differences. Sex is a proxy for multiple genetic, developmental, and endocrine influences that vary across individual mammals [15–17] and which may be expected to influence how complex, multidimensional environments are sampled and experienced [18–20]. Indeed, sex differences (and gender differences in humans) have been shown in a variety of simple value-based decision-making tasks, but these differences are not always consistent, suggesting the presence of latent variables other than straightforward reinforcement learning properties that may influence how individuals strategically sample their environment and thus learn [21–23]. However, much of the work identifying differences in performance across males and females has used tasks with low trial counts, choices that vary on only one dimension, and/or confounded rewarding and punishing outcomes that are not well-suited to computationally elucidating the strategies employed during choices. As a result, whether sex differences in value-guided decision making reflect sources of diversity in the strategies employed in sampling multidimensional environments is unknown.

To identify potential sex differences in the strategies employed during value-based decision making, we examined male and female mice as they performed a two-dimensional decision-making task: a two-armed visual bandit[1,2,4,24–28]. While all animals eventually reached the same performance level, female mice learned more rapidly than males and acquired more rewards over the course of learning. Because choice could vary in multiple dimensions in this task[29,30], we were able to consider the possibility that individual animals were adopting different strategies. We found that sex explained a substantial fraction of the individual variability in strategy. Female mice used a systematic strategy where they preferred to choose options in one spatial location, constraining the decision-space and accelerating their learning about image values, even though rewards were counterbalanced across spatial location. Conversely, males simultaneously considered information from both image and spatial dimensions simultaneously, were highly sensitive to the stochastic experience of reward, and changed their own choice strategies frequently. During early learning, gene expression for the neuronal activation marker c-fos in the nucleus accumbens and prefrontal cortex significantly correlated with the extent to which female animals (but not males) used a systematic strategy. These results show that individuals can adopt widely divergent strategies for interacting with the same uncertain world, and that sex is a substantial factor in guiding these strategies.

## Results

Age-matched male and female wildtype mice (n=32, 16 per sex, strain B6129SF1/J) were trained to perform a visually-cued two-armed bandit task in touchscreen operant chambers (**Figure 1a**). This task design was similar to those employed in humans and nonhuman primates [1,2,4,24– 28,31,32], in contrast to the spatial bandit designs frequently employed with rodents [33–36]. Animals were presented with a repeating set of two different image cues which were associated with different probabilistic reward outcomes (**Figure 1b**).The reward schedule (80%/20%) was held constant throughout the session. In contrast with spatial bandit designs, here reward contingencies were yoked to image identity, which was randomized with respect to location in the chamber on each trial. This means that the sides (left/right) where image cues appeared were not informative of the reward contingencies. We repeated the task with six different sets of image pairs. Two out of the six tested image pairs were excluded from the study due to extremely high initial preference (>70%) for one image over another. We included four images pairs with equal initial preference for each image and quantified behavioral data in bins of 150 trials for each animal.

**Figure 1.**
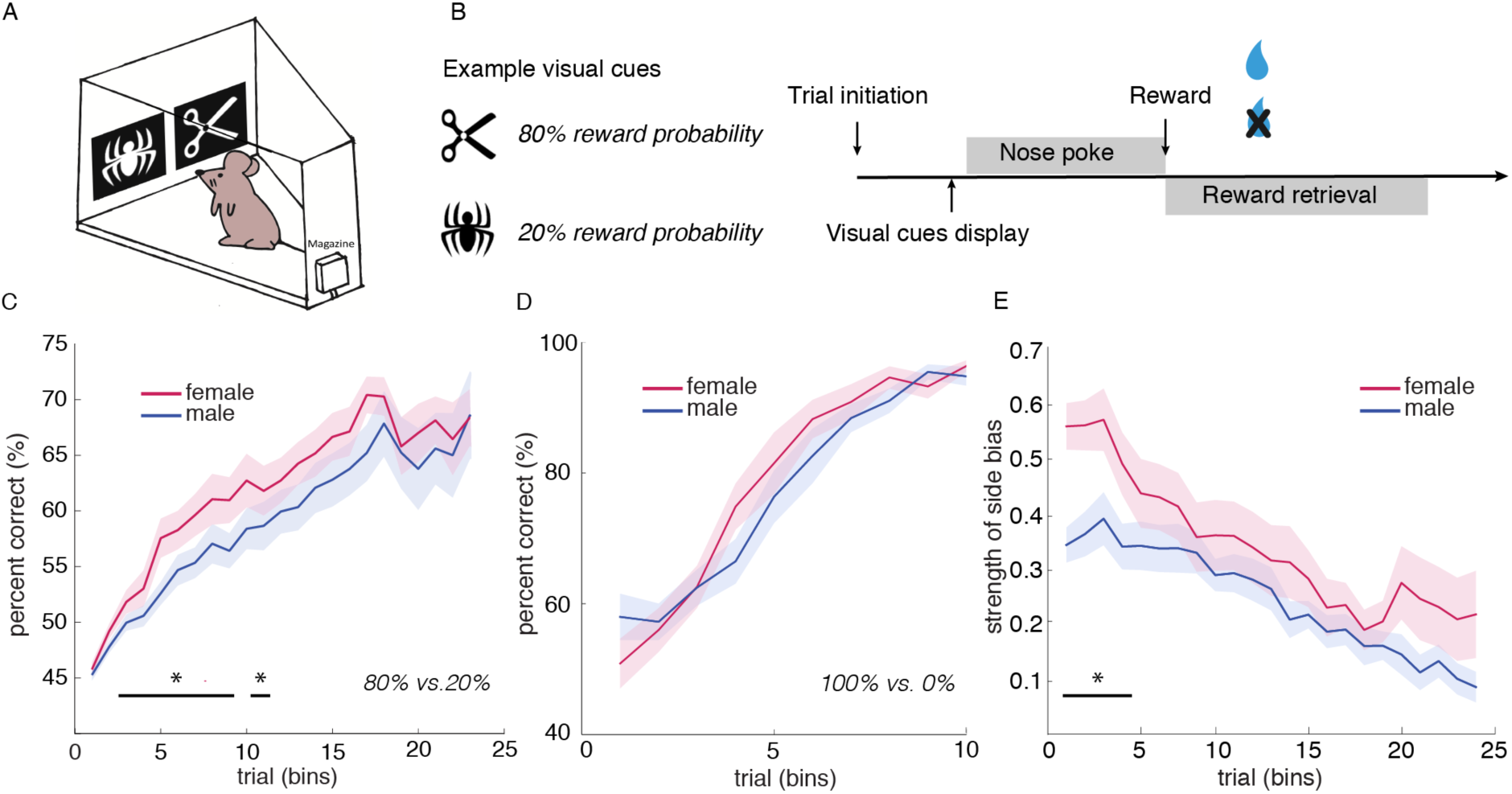
Females showed accelerated acquisition of the high reward probability image in a stochastic two-armed visual bandit task. A) Schematic of the mouse touch-screen operant chamber used in our task. B) Schematic of two-armed Visual Bandit task. Images varied between the two locations across trials. The reward probabilities for two images are 80% and 20%, respectively. C) Average learning performance (percent correct) across four repetitions of the task with four pairs of images. While both males and females reached the same final performance, females displayed an accelerated learning curve. D) No sex difference in learning performance was observed in deterministic reward schedule (100%/0%). E) Females displayed stronger side bias for item selection on the touchscreen, regardless of the direction of lateralization, early on in learning. This behavior lateralization disappeared as female mice learned the task. Data shown as bins of 150 trials. * indicates p < 0.05. Graphs depict mean ± SEM.

### Females showed accelerated reinforcement learning, but males and females reached equivalent final performance

To examine learning performance, we calculated in bins of 150 trials the average percentage of choosing the high-value image (23 bins in total). Regardless of sex, mice were capable of eventually learning which image was associated with the higher reward probability (**Figure 1c**, GLM, main effect of sex, p = 0.51, β1 = - 0.05; main effect of number of trials, p < 0.0001, β2 = 0.10, see equation 1 in Methods). However, we repeatedly observed that females learned the image pair discrimination significantly faster than did males (GLM, interaction term, p < 0.05, β3 = −0.02). We compared these results to a deterministic version of the task in the same animals, in which one image was always rewarded (100%) and the other was never rewarded (0%). We did not find any significant sex difference in rate of learning across trials in the deterministic task (**Figure 1d**, GLM, interaction term, p = 0.38, β1= −0.004, see equation 1 in Methods), suggesting the difference was revealed by the stochastic experience of reward.

To determine the origins of this sex difference in early performance, we first considered that sex differences in early learning might reflect differences in the rate of value-updating and/or the level of random noise in choice--the typical parameters of reinforcement learning models. We fit a delta-rule reinforcement model [4,33,37,38] to measure individual differences in the learning rate parameter and noise parameter, based on choices of images. However, the likelihood surface of the model given by parameters learning rate (α) and inverse temperature (β) was flat, which prevented parameter optimization. This suggested to us that the basic RL model based on images as the sole choice dimension could not characterize individual differences in learning in this task.

Since rodents are generally highly spatial, we hypothesized that mice might have a bias towards using spatial information earlier in the task, before switching to use image information, as demonstrated by the high rates of reward late in training. Consistent with our hypothesis, we observed a short period of high side bias in females early in learning (**Figure 1e**), which could include a preference for either the left or right side and seemed to precede the acquisition of the reward contingency. Following the decrease of side bias, female mice improved their percentage of choosing high-value image. (GLM, main effect of sex, p < 0.001, β1= −0.129; main effect of number of trials, p < 0.001, β2 = −0.017, see equation 1 in methods). Strength of side bias independent of left or right side was calculated using methods described in previous behavioral lateralization literature [39].

### Females systematically reduced the dimensions of the task by strongly preferring one side

A side bias is only one of several local strategies that mice could have been using as they learned the reward contingencies in the task. For example, another local strategy is a spatial win-stay strategy, where the side of the last choice is repeated if and only if it was rewarded. Alternatively, an image win-stay strategy would repeat the last image, if and only if it was rewarded, or an image bias strategy would simply select the previous image, regardless of reward. To understand how different animals employed these different local strategies and processed through them over time, we constructed a generalized linear model (GLM) to predict each choice, based on a weighted combination of these local strategies. The model had a term to account for two classes of basic strategies: outcome-independent strategies and outcome-dependent, win-stay, strategies (**Figure 2a**, see Equation 4 in methods). Outcome-independent strategies (image repeat and spatial repeat) captured the tendency of repeating either the side or the image of the previous trial, regardless of the outcome. Conversely, outcome-dependent strategies followed a win-stay lose-shift policy for either a side or an image, which captured the tendency to only repeat the side or the image of the previous trial when it was rewarded. Fitting the GLM allowed us to estimate how much each of these four strategies was employed within each animal on each bin of trials. We will call this set of weights--the precise pattern of local strategies employed over time--the “global strategy” employed by each individual animal.

**Figure 2.**
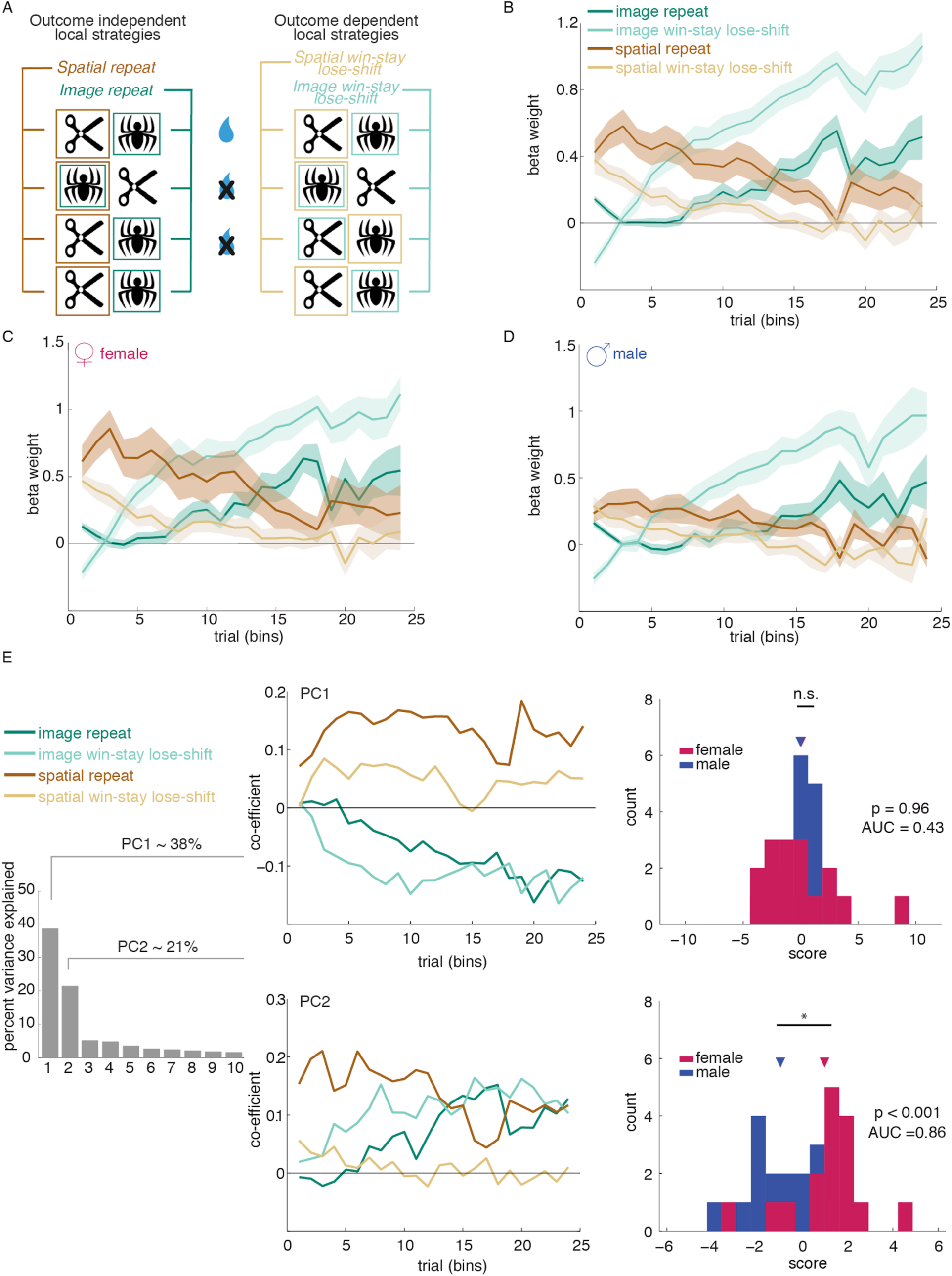
Female mice use a procession of strategies, initially using a spatial bias followed by a switch to responding based on image domain. A) Schematic of four basic local strategies based on choice and reward history of image and spatial dimensions of the task. B) A generalized logistic regression model revealed a global strategy - a clear procession of four local strategies - that mice started repeating one side before switching to choosing the reinforced image. C) female mice displayed more pronounced global strategy procession from spatial-based strategy to image outcome-based strategy. D) male mice displayed increased image win-stay lose-shift over time but no prominent global strategy in the early learning stage was observed. E) A principal component analysis (PCA) was conducted on the estimates of global strategy strength over time across all animals regardless of sex. Principal component (PC) 1 and 2 accounted for about 60% of the variance. PC 1 described a general preference for responding based on image value and did not differ between sexes. PC 2 captured the same global strategy procession reflected in the generalized logistic model - a strong contribution of the “side repeat” behavior early in training, followed by a rapid transition to “image outcome”, indicative of a sudden shift away from “where” and towards “what” in solving the task. Projecting each animal onto this PC 2 showed a clear separation between the sexes (blue male, pink female), AUC = 0.86, p < 0.001. This suggests that the strategy procession from spatial repeat to image outcome is a female-specific strategy. Note that a few males are positive for Principal Component 2, and their individual behavior supports that these males also employed this strategy to a weaker extent. In contrast, the few females that are negative for this Principal Component did not show evidence of having learned the task. Data shown as bins of 150 trials. Graphs depict mean ± SEM.

Across all animals, we found a global strategy where a specific procession through local strategies was used when learning image pairs (**Figure 2b**). First, animals showed an early tendency towards outcome-independent spatial repeat, giving way to a later interaction between reward outcome and image choice, with outcome-insensitive image repeat (the optimal strategy) increasing in the later stages of testing. To examine whether sex influenced the strength of this strategy procession, we compared the global strategy used by male and female animals. We observed a consistent and pronounced pattern of strategy procession only in females (**Figure 2c**). In contrast, in males, the weight of both image-based strategies simply increased slowly over time (**Figure 2d**), with little change in spatial strategies. Thus, neither a procession of multiple strategies nor a prominent strategy in the early learning stage was observed in male mice.

The sex difference in this strategy selection was intriguing, but it could have been driven by only a few females. Therefore, we next characterized individual variability in strategy via principal component analysis. Here, we estimated the major axes of interindividual variability in strategy, meaning in the unique combinations of the four strategy weight vectors over time and across all animals, regardless of sex. Principal components (PC) 1 and 2 captured the majority of the interindividual variance: 59% of the variability between animals (**Figure 2e**). PC1 reflected a global preference for a side or an image and did not significantly differ between sexes (p > 0.9, AUC = 0.43). The mean principal component scores of PC1 for females and males were 0.03 and −0.03, respectively. The mean PC score difference between females and males (mean(F-M)) was 0.07 (95% CI = [−1.70, 1.80], t(30) = 0.08). Critically, PC2 mirrored the same procession of strategies observed in female, but not male mice (**Figure 2c-d**). This suggests that the extent to which individuals used this procession of strategies explained a large fraction (22%) of the interindividual variability in our animals. The mean PC score of PC2 for females was 0.98 and for males was −0.98. The mean PC score difference between females and males (mean(_F-M_)) was 1.96 (95% CI = [0.87, 3.05], t(30) = 3.67). Moreover, knowing the PC2 score (the projection of an individual animal’s behavior onto PC2) allowed us to discriminate male and female animals with remarkable accuracy (reciever operating characteristic analysis, AUC = 0.86, significant discrimination: p < 0.001). No otherPCs differed between sexes (p > 0.4, AUC < 0.6). Together, these results suggest that the choice of what strategy to follow explains substantial individual variability in multidimensional decision-making, and that differences in strategy can depend on sex.

There are two possible explanations why females consistently implemented the strategy procession captured by PC2. One hypothesis is that this early side bias reflected an *energy saving* strategy that saved time and/or effort by just repeating the same side. Alternatively, this early side bias could be an *cognitively effortful* strategy to constrain decision-making to one dimension. These two views make different predictions of the relationship between the use of strategy procession captured by PC2 and reaction times (RTs), which were defined as the time between the onset of two visual stimuli and the registration of a nose poke response on one of the two stimuli. If side bias was an effort saving strategy, then the animals who score highest on PC2 should also make the fastest decisions. On the other hand, if side bias was a cognitively effortful strategy, the speed of decision-making should be slower in animals who use this strategy. To test these two hypotheses, correlation analyses were run to assess the relationship between the use of PC2 strategy and reaction time. PC2 scores were positively correlated with reaction time (**Figure 3a**, Spearman’s correlation, r_s_ = 0.452, p = 0.009; Pearson’s correlation, r = 0.347, p = 0.051), suggesting that the animals who used the early side bias strategy made slower decisions. This suggests that this strategy is effortful, rather than energy saving. There were no significant correlative relationships between reaction time and other PCs.

**Figure 3.**
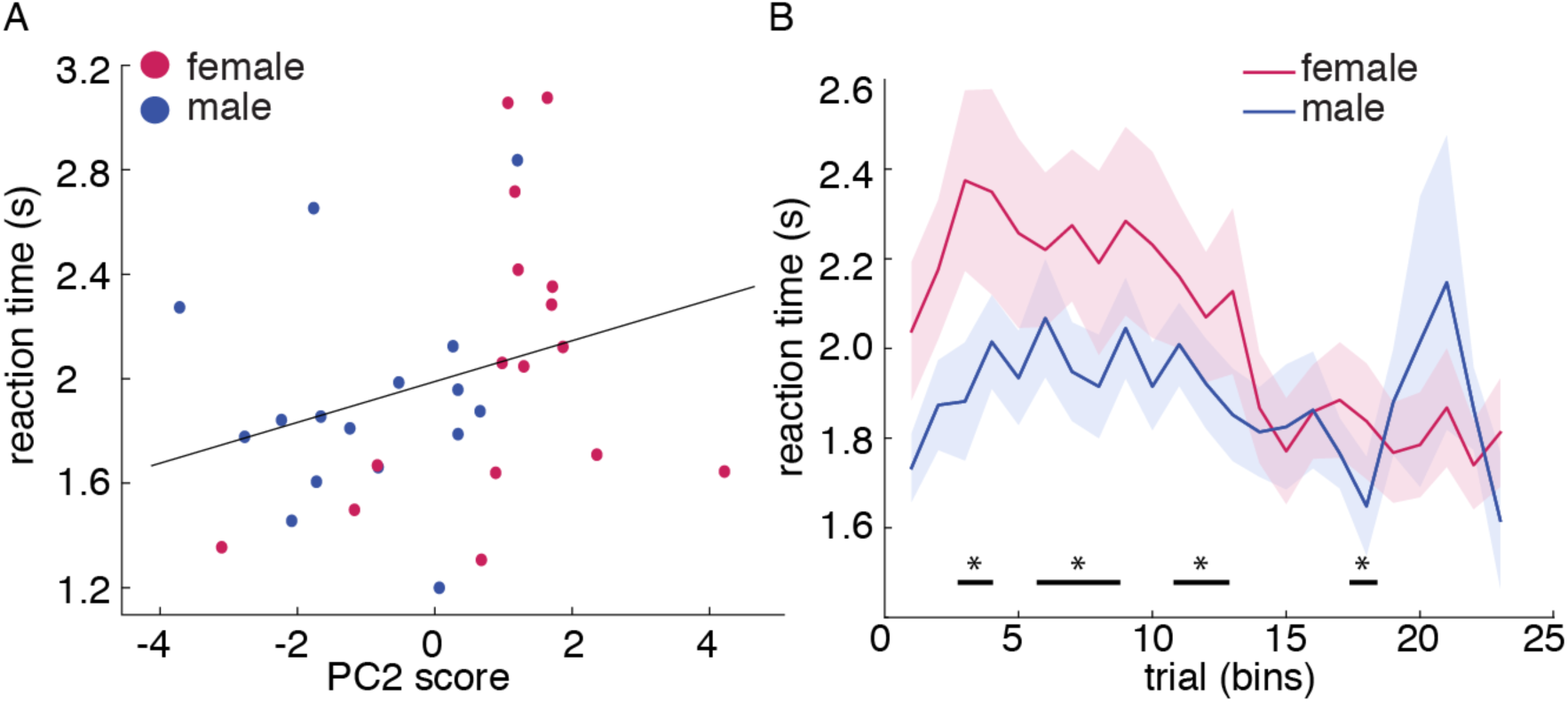
The side-to-image strategy procession captured by Principal Component (PC) 2 is a cognitively demanding strategy but not a time-saving strategy. A) Correlation analyses revealed a significant positive correlation between PC2 scores and reaction time. The decision-making time was longer within animals primarily used PC2 strategy. B) Predominantly using PC2, females responded slower during early learning (GLM, interaction term, β3 = 0.03, p = 0.0007). Note that, the slow reaction time during early learning in females matches with the time period (bin 1-10) during which females relied on side-bias “heuristics” for decision-making. Data shown as bins of 150 trials. * indicates p < 0.05. Graph B depicts mean ± SEM.

If the side bias in females was a cognitively effortful strategy, we would expect to see that female mice are slowest at making their choices when they are most engaging this strategy: during early learning. To test this hypothesis, we computed average RTs across 23 bins of 150 trials for males and females. Consistent with our hypothesis, females responded slower during early learning (bin 1-15) (GLM, interaction term, β3 = 0.03, p = 0.0007) and significantly slower than males across all trials (GLM, main effect of sex, β1 = −0.62, p < 0.0001). The mean RT across all trials for males was 1.89 seconds with standard deviation of 0.13, and the group average for females was 2.04 seconds with standard deviation of 0.21. The reaction time decreased as the animals ran more trials in both males and females (**Figure 3b**, GLM, main effect of number of trials, β2 = −0.04, p<0.0001, see equation 1 in methods). Thus, female mice were slowest during the period in which they were using the side bias strategy the most, again consistent with the idea that this is a cognitively effortful strategy, rather than an energy saving one.

### Males varied strategies over time in response to immediate past reward

Although our analyses captured the procession of strategies employed across essentially all female mice that learned the task, they provided little insight into what the males were doing. One likely explanation is that males were more inconsistent than females. Males could be more inconsistent than females for any one of three reasons: hypothesis 1) males were more *random* (and thus each choice would be unpredictable within an animal), hypothesis 2) males were more *idiosyncratic* and less uniform as a group (and thus responses would differ across individuals, but still be predictable within an individual), or hypothesis 3) that males were more *changeable* (and thus a given male was predictable in the sense that he was not random, but he was still more likely to change his strategic approach from one epoch to the next).

To test hypothesis 1 (randomness), we asked whether male choices were more or less predictable than female choices. We reasoned that if males were just choosing randomly, it would be impossible to predict their choices. Therefore, within each block of trials in each animal, we calculated conditional mutual information [40,41], which quantifies the dependence of current choices (side, image) on the choice of previous trial, given the outcome of the previous trial. If the current choice an animal made was random, it would be independent of the choice and outcome of the previous trial, and we would expect to see low mutual information, shown as uniform “bands” on the probability heatmap (**Figure 4a**). Conversely, if the current choice was heavily influenced by the content of the previous trial, we would expect to see high mutual information, shown in a more checkered or selective pattern on the probability heatmap. We calculated conditional mutual information for each trial bin across sexes. We found that mutual information decreased over time in both sexes, reflecting the gradual acquisition of the strategy of choosing high reward probability cue regardless of the outcome of the previous trial (**Figure 4a**, GLM, main effect of number of trials, β2 = −0.001, p =0.0002, see equation 1&5 in Methods). However, surprisingly, the mutual information of male mice was *higher* than that of females (main effect of sex, β1 = 0.043, p < 0.0001), particularly early in learning (interaction term, β3 = - 0.002, p < 0.0001). Thus, males were, if anything, less random than females.

**Figure 4.**
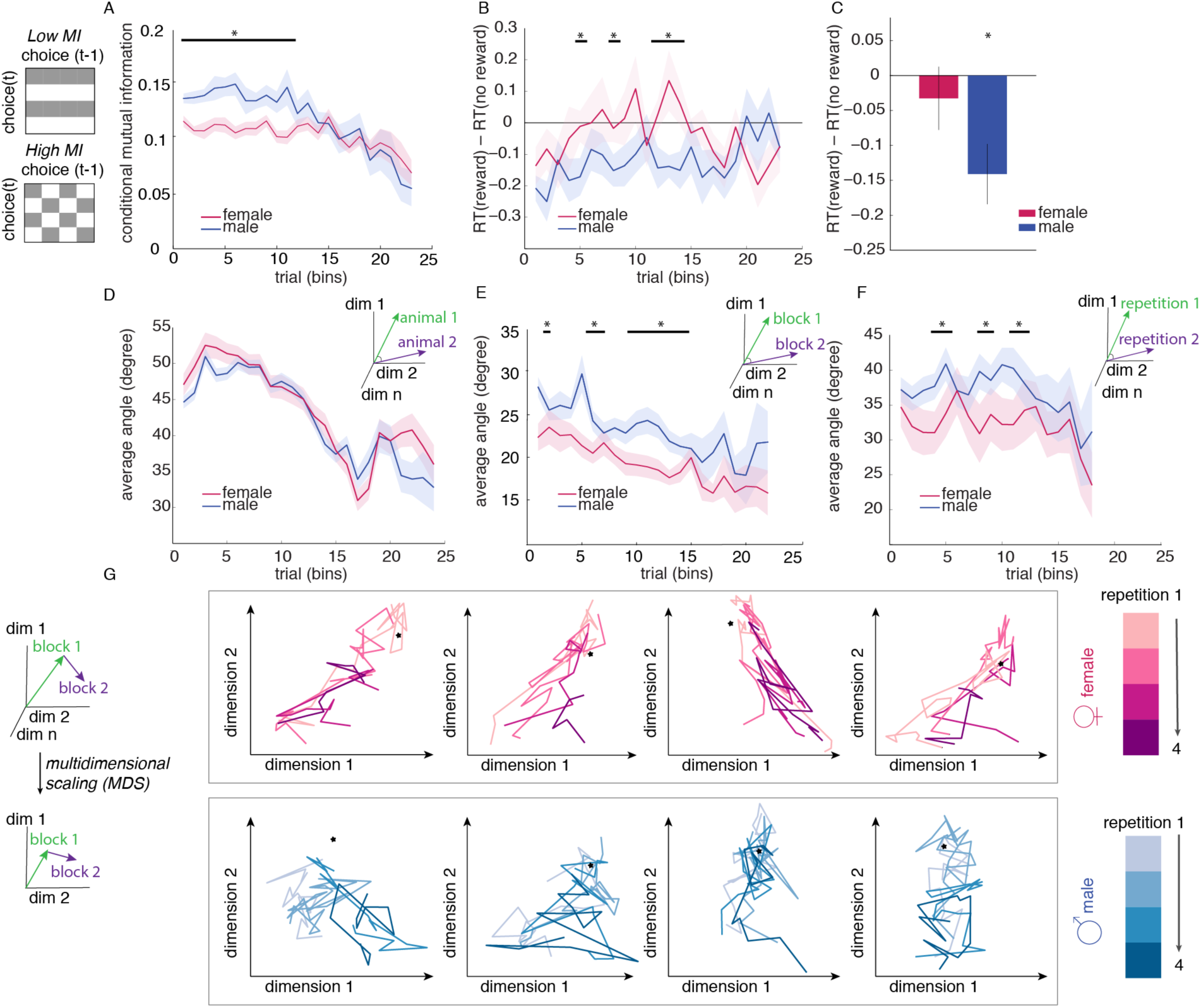
Male mice were more likely to differ from themselves over time, with choice patterns dependent on past outcomes. We expressed each animal’s history-dependent choice pattern as an 32-dimensional vector of joint probabilities and measured the angle between vectors, which is proportional to the step between them on a strategy simplex. A) Illustration of choice patterns of low mutual information and high mutual information. If choice on trial *t* is independent of choice on the previous trial (*t-1*), probability heatmap should show band-like pattern (choosing the same choice regardless of the previous choice). Conversely, high mutual information has more checkered choice patterns. Conditioned mutual information is higher in males, indicating that responses are more uniquely affected by the previous trial variables than they are in females. B) One-sample t-test was conducted across bins to compare the difference in reaction time (RT) between rewarded and unrewarded trials to 0 (when there is no effect of past outcome on the reaction time). Male mice have significant RT effects on the last reward. There was no difference in reaction time between rewarded and unrewarded trials in female mice. The bins marked by asterisks have p < 0.05 for the one-sample t-test. C) average RT effect of last reward across all trials. Overall, male responded faster when the last trial was rewarded than unrewarded. D). Males and females were equally variable between animals within sex. An individual male is no more different from other males in behavior than a female is from other females. E) Choice patterns of a given male compared to himself over bins of 150 trials were more variable than in a given female compared to herself. F) Choice patterns of a given male to himself were more variable and divergent across repetitions of the same task than in females compared to themselves across repetitions. G) Multidimensional scaling (MDS) was used to visualize animal’s strategy path across trials and repetitions by reducing the dimensionality of the strategy space. Each of the four colors within one sex represents one repetition of the task. The star represents the point of optimal strategy for this task, which is to choose the high reward probability image. In both males and females, the strategy paths showed a gradual approach to the optimal strategy point, indicating that both sexes were able to learn the optimal strategy. The strategy paths of females are consistent and similar across repetitions, suggesting that female mice used a similar strategy every time to solve the problem. On the other hand, in male mice, the steps between each bins of 150 trials in the strategy space were larger, suggesting higher variability in choice patterns. Thus, male mice used divergent strategies throughout learning and used different approaches each time to learn the same task. Data shown as bins of 150 trials. * indicates p < 0.05. Graphs depict mean ± SEM.

One possibility is that the high mutual information in males was a result of increased sensitivity to the last reward, meaning that the last reward had a bigger effect on the males’ idiosyncratic decisions about what to do next. To estimate sensitivity to reward, we examined how reward outcomes affected the reaction time (RT) on the next trial in both males and females. If the decision of an animal was not affected by the outcome of the past trial, then we would expect to see no difference between reaction time for last rewarded and last unrewarded trial (RT_reward_ – RT_no reward_ = 0). Males responded significantly faster when they had just received a reward from the previous trial (**Figure 4b and 4c**, one-sample t-test, mean RT effect = −0.14, 95% CI = [− 0.23, −0.05], t(15) = −3.38, p = 0.004). Conversely, the reaction times of females were not affected by the outcome of the last trial (one-sample t-test, mean RT effect = −0.03, 95% CI = [− 0.13, 0.06], t(15) = −0.75, p = 0.47). These results again suggested that female decisions were not affected on a trial-by-trial basis by the outcome of each trial because they followed a global strategy while male choices that were heavily affected by recent rewards.

Although males were, if anything, less random in their decision-making than females, it remained possible that the apparent lack of local strategies occurred because male strategies were inconsistent--either because of idiosyncratic differences between males (hypothesis 2) or changeability within males (hypothesis 3). To do this, we developed a technique for comparing how similar one set of choices was to other set of choices. We expressed the choices in each bin as a probability vector, with each element of the vector reflecting the probability of that unique combination of behaviors {last choice, last outcome, this choice}. The average angle between any two of probability vectors across animals, trial bins, or image pairs is then a measure of the variability in choices between those two conditions. Males were not more idiosyncratic than females on a population level; that is, the choices of any male were not more variable from other males than any female’s choices were from other females (**Figure 4d**, GLM, main effect of sex, β1 = −1.47, p = 0.11, see equation 1 in Methods). However, the males were more variable *within themselves*, both across bins within one image pair (**Figure 4e**, GLM, main effect of sex, β1 = 4.24, p < 0.0001, see equation 1 in Methods) and across multiple image pairs (**Figure 4f**, GLM, main effect of sex, β1 = 4.54, p = 0.047, see equation 1 in Methods). Overall, the variability in choices decreased across time, as the divergent strategies used by individual animals started to converge to the optimal strategy in this task, which is to choose the high-value image consistently (GLM, main effect of number of trials, within sex between suject: β2 = −0.78, p < 0.0001; within subject between bins: β2 = −0.359, p < 0.0001). Together, these results suggest that individual males displayed divergent choice patterns and were changing between complex strategies over time and the repetition of the same task, while females largely adopted a shared, systematic approach to learning.

To visualize animals’ patterns of choices expressed in the probability vectors, we used multidimensional scaling (MDS) [42–44] to reduce the dimensionality of strategy space, allowing us to project the high-dimensional “strategy path” throughout learning onto a 2 dimensional space. This allows us to easily visualize the similarity between patterns of choice across animals over time and across repetitions. Each color path represents a strategy path used in a different repetition of the task (4 repetitions in total). **Figure 4g** shows examples of strategy paths of males and females. The optimal strategy in this bandit task, which is to choose the high value image consistently regardless of the outcome, is represented by a star in the low dimensional space. Both males and females “strategy path” showed gradual approximation to the optimal strategy over time. Consistent with the quantification described above, the strategy path of males are visibly more variable and different across repetitions of the task, whereas the strategy path of females were more unified and consistent across repetitions.

### Sex mediated the ability of neuronal activity to explain strategy selection

The ability to learn and perform bandit tasks is highly sensitive to alterations in corticolimbic structures. However, it remains unclear how alterations in these structures predict choice strategy, much less sex differences in choice strategy. To address this question, we examined neuronal activity in several corticolimbic brain regions by examining the expression of *c-fos*, an immediate early gene often used as a marker of neuronal activation. The animals from the previous figures were sacrificed after the second day of a new, final image-reward pairing (each animal having completed 400-500 total trials), corresponding to when the female side bias was greatest. We compared mRNA expression level for *c-fos* across five brain regions, including nucleus accumbens (NAc), dorsal medial striatum (DMS), amygdala (AMY), hippocampus (HPC), and prefrontal cortex (PFC), using quantitative real-time PCR (**Figure 5a**). In each of the five brain regions, females had a higher c-fos expression level than did males (unpaired t-test, NAc: t(30) = 2.41, p = 0.02; DMS: t(30) = 2.31, p = 0.03; AMY: t(30) = 4.05, p < 0.001; HPC: t(30) = 2.74, p = 0.01; PFC: t(29) = 3.163, p = 0.003).

**Figure 5.**
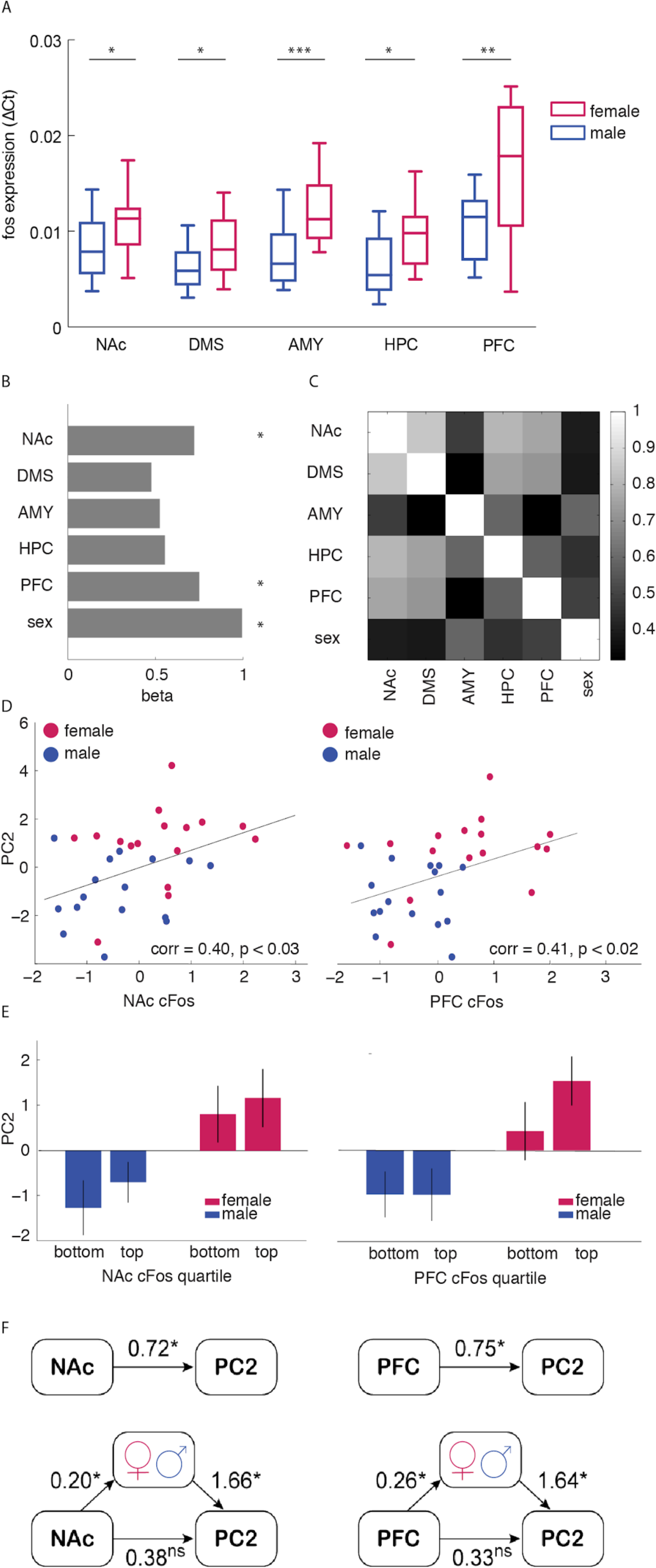
Both sex and neuronal activity can account for strategy selection, but sex mediated the ability of neural activity to explain strategy selection. **A**) cFos gene expression (qRT-PCR) in five brain regions: nucleus accumbens (NAc), dorsal medial striatum (DMS), amygdala (AMY), hippocampus (HPC), and prefrontal cortex (PFC). Female mice showed elevated c-fos expression across all five brain regions. Asterisks marked significant difference between sexes (*: p < 0.05 **: p < 0.01 ***: p<0.001). **B**) Heatmap of correlation matrix of c-fos expression level among five brain regions. **C**) c-fos expression in NAc and PFC, and sex, predict the use of PC2 strategy (GLM, NAc: β1 = 0.72, p = 0.02; PFC: β5 = 0.75, p = 0.02; sex: β6 = 0.99, p = 0.0009). Asterisks marked significant beta weights (p < 0.05). **D**) cFos gene expression in NAc and PFC is significantly correlated with the weight of PC2. **E**) The use of PC2 strategies procession was analyzed with a 2 (sex: male versus female) x 2 (c-fos expression quartile in NAc/PFC: bottom versus top) between-subjects ANOVA. The main effect of sex was significant for both NAc and PFC (NAc: F (1,28) = 12.87, p = 0.001; PFC: F (1,28) = 13.47, p = 0.001). F) Causal modeling of the relationship between gene expression level in NAc and PFC and the weight of PC2. The models on top are direct models, indicating that c-fos expression levels in both NAc and PFC are significant predictors of the weights of PC2. The bottom models are the mediation models, in which sex mediated the relationship between neural activity (c-fos expressions in NAc and PFC) and strategy selection (weights of PC2). The arrows are regressions. Paths are labeled with estimated coefficients and significant coefficients are marked by asterisks. The strength of the direct model is greatly reduced and became non-significant when accounted for the mediating effect of sex. This suggests that sex mediated neural measures in explaining strategy selection. Graphs depict mean ± SEM.

To understand whether activation of any of these brain regions correlated with the side bias strategy, we constructed a GLM to predict PC2 from c-fos expression level in each brain region and sex. The results suggested that only two regions, the NAc and PFC predicted strategy use, as indexed by PC2 score (**Figure 5b**, GLM, NAc: β1 = 0.72, p = 0.02; DMS: β2 = 0.48, p = 0.14; AMY: β3 = 0.52, p = 0.10; HPC: β4 = 0.55, p = 0.08; PFC: β5 = 0.75, p = 0.02; sex was included as a variable in the model and was also significant: β6 = 0.99, p = 0.0009, see equation 2 in Methods). Because each region was also correlated with sex to differing extents (and sex independently predicted PC2), we next asked whether NAc and PFC were the best predictors of PC2 because these regions were the most strongly correlated with sex (**Figure 5c**). However, the predictive effect of NAc and PFC c-fos expression was not because NAc and PFC were the most highly with sex. Instead, sex was most strongly correlated with AMY, which was not a significant predictor of PC2. To confirm that these correlations between regional activation and early side bias strategy was meaningful, we fit the same GLM to predict PC1, and none of the predictor variables were significant. We confirmed these results with a Pearson product-moment correlation coefficient, which again suggested a significant positive correlation between c-fos expression in NAc/PFC and PC2 scores (**Figure 5d**, NAc: r = 0.40, n = 32, p < 0.03; PFC: r = 0.41, n = 32, p < 0.02; averages across a median split of PC2 within each sex are illustrated in **Figure 5e**; main effect sex: NAc: F (1,28) = 12.87, p = 0.001; PFC: F (1,28) = 13.47, p = 0.001).

Next, we asked whether an animals’ sex altered the relationship between NAc and PFC c-fos activity PC2 scores. To do this, we used a structural equation modeling (SEM) approach [45,46] to analyze the structural relationship between sex, gene expression, and PC2 and latent constructs (**Figure 5f**). First we used a direct model and regressed c-fos expression of either NAc or PFC on strategy selection, both NAc and PFC were significant direct predictors of PC2 scores (NAc: β = 0.72, p = 0.022; PFC: β = 0.75, p = 0.019, see equation 6 in Methods). Then we fit a mediation model that allows us to understand how sex influences neural activation in NAc and PFC, which in turn influences strategy selection. Regressing the mediator variable sex on c-fos expression in NAc/PFC confirmed that neural activation is a significant predicor of the mediator sex (NAc: α = 0.20, p = 0.024; PFC: α = 0.26, p < 0.004, see equation 7 in Methods). When we regressed strategy selection on both the mediator variable (sex) and independent variable (neural activation in NAc/PFC), the result showed that the mediator sex was a significant predictor of strategy selection (NAc: β1 = 1.66, p = 0.008; PFC: β1 = 1.64, p = 0.012), and the strength of the direct model is now greatly reduced and became non-significant when accounted for the mediating effect of sex (NAc: β’ = 0.38, p = 0.20; PFC: β’ = 0.33, p = 0.31). The Sobel (1982) first-order test was used to assess the presence of mediation [46]. The indirect effect was calculated as the product of coefficients and was significant for both NAc and PFC (NAc: αβ’ = 0.34, z = 1.836, p < 0.039; PFC: αβ’ = 0.42, z = 2.035, p < 0.026). Together, these results suggest that the relationship between PFC and NAc c-fos and PC2 differed, depending on the animals’ sex. This suggests that sex-linked mechanisms gate the relationship between these circuits and strategic decision-making and highlight these regions as promising targets for future studies looking at the effects of sex on the neural circuits responsible for implementing strategic learning.

## Discussion

By training male and female mice on a stochastic two-dimensional decision-making task, we were able to evoke a range of problem solving strategies across individuals. In this task, each cue has two dimensions - the identity of the image and the location of the image. Animals had to explore the reward value associated with both cue dimensions to determine which were most predictive of reward. Although both male and female mice eventually learned the right strategy, choosing the high-value image, female mice learned faster. The richness of this task allowed us to uncover sex differences in *how* the animals achieved the associations across time. We discovered that female mice adopted a consistent and systematic approach where they processed through different strategies over time. Early in learning, they constrained their search space by only sampling the outcomes of images on one side (left or right). This approach, which occurred when animals were most uncertain about the best choice, reduced the number of dimensions they were learning about and permitted more rapid acquisition of the image-value association. In contrast, males employed a strategy of decisions that seemed to combine both image and spatial location, changed frequently, and was strongly influenced by the immediate prior experience of reinforcement. While both sexes eventually reached equivalent levels of performance, our data reveal that the strategic paths journeys individual animals take to get there can vary dramatically, implicating the potential for wide divergence in neural circuit mechanisms in normative decision making.

Sequential decision-making and learning in rodents is often studied with spatial bandit tasks, in which reward probabilities are linked to left and right levers or sides that are visually identical [1,4,11,13,33,47]. In these spatial bandit tasks, side bias in choice has sometimes been reported as a behavioral artifact and animals displaying such bias were often excluded from experiments [48–50]. However, in the current task, both the side and the identity of the image cues could have been informative of the reward probabilities (although they were not). In principle, animals could simultaneously sample both dimensions to learn side values and image values at the same time. However, in practice, it appears that the early side bias in female mice “jump-started” their learning by controlling for space while exploring choice-outcome values of the images, which in this task happened to be the more informative dimension of reward. Intriguingly, this suggests that females were covertly learning about the correct cue dimension while behaviorally selecting the wrong item, and were able to convert this to successful learning due to the stability of the task structure. This view suggests that we should be able to design tasks that prevent the successful use of this strategy, and which might therefore shift the presentation of the sex difference in decision-making.

Our data implicate the prefrontal cortex (PFC) and nucleus accumbens (NAc), part of the ventral striatum, in the differences in strategy between males and females. These regions have been widely implicated in reward-guided decision making, but so have the other regions we tested for which we didn’t find a significant relationship to behavior [2,29,51]. One possibility is that the PFC and accumbens are particularly engaged in strategic decision-making. This resonates with previous studies that have implicated the PFC in implementing strategies and rule-guided behaviors [47,52–57] and the NAc in selecting and implementing learning strategies [29,30]. Implementing different strategies produces changes in how different choice dimensions are represented in the PFC and NAc [58], and lesions in the NAc can drive animals towards a low-dimensional action-based strategy or prevent animals from switching between strategies [2,30]. The PFC is also sensitive to gonadal hormones during risky decision making [59], and dopaminergic function in the accumbens regulates risky decision making in a sex-specific manner [60], perhaps due to sex differences in dopamine neurons [61]. Here, the relationship between both PFC and NAc and strategy use was mediated by sex, suggesting that whatever the relationship between these regions and strategic decision-making, it is likely to be sex-biased.

One fundamental unanswered question is *why* females as a group employed a highly similar and consistent strategy. Zador (2019) recently proposed that much of animal behavior is not dictated by supervised or unsupervised learning algorithms, but are largely shaped by biological constraints [62]. Energy-conserving and habitual behaviors are prevalent in female animals, including during foraging [22,63,64]. The biological constraints and organization imposed by the multiple mechanisms of sexual differentiation are known to drive a tuning of the circuits important for reward-guided decisions [61,65]. Exogenously applied gonadal steroid hormones are known to modulate cost/benefit decision-making. While testosterone promotes effort expenditure and impulsive behavior [66–68], estradiol reduces high-effort choices [69–71]. In addition, sex chromosomes exert independent influences on reward-guided behaviors [15] with elevated habit in XX carriers and increased effort in XY carriers, regardless of gonadal hormone status [72,73]. Of course, these impacts of sexual differentiation are graded, rather than dichotomous across the sexes. Indeed, here we found that a small number of males showed some tendency to use the female strategy, implicating graded mechanisms of early-life and/or pubertal masculinization in the degree of adoption of sex-biased exploratory strategies we observed here. An intriguing possibility is that the unified, consistent, and systematic strategy we observed in female mice, as well as the volatile and diverse strategy we observed in male mice in the same task, may emerge from evolved sex-biased strategies for foraging in the wild that were critical to survival for the species as a whole.

## Methods

### Animals

Thirty-two BL6129SF1/J mice (16 males and 16 females) were obtained from Jackson Laboratories (stock #101043). Mice arrived at the lab at 7 weeks of age, and were housed in groups of four with ad libitum access to water while being mildly food restricted (85-95% of free feeding weight) for the experiment. Animals engaging in operant testing were housed in a 0900– 2100 hours reversed light cycle to permit testing during the dark period, between 09:00 am and 5:00 pm. Before operant chamber training, animals were food restricted to 85%-90% of free feeding body weight and had been pre-exposed to the reinforcer (Ensure). Pre-exposure to the reinforcer occured by providing an additional water bottle containing Ensure for 24 hours in the home cage and verifying consumption by all cagemates. Operant testing occurred five days per week (Monday-Friday), and the animals were fed after training with ad lib food access provided on Fridays. All animals were cared for according to the guidelines of the National Institution of Health and the University of Minnesota.

### Apparatus

Sixteen identical triangular touchscreen operant chambers (Lafayette Instrument Co., Lafayette, IN) were used for training and testing. Two walls black were acrylic plastic. The third wall housed the touchscreen and was positioned directly opposite the magazine. The magazine provided liquid reinforcer (Ensure) delivered by a peristaltic pump, typically 7ul (280 ms pump duration). ABET-II software (Lafayette Instrument Co., Lafayette, IN) was used to program operant schedules and to analyze all data from training and testing.

### Operant Training

#### Pretraining

animals were exposed daily to a 30-min session of initial touch training, during which a blank white square (cue) was presented on one side of the touchscreen, counterbalancing left and right between trials. This schedule provided free reinforcement every 30 seconds, during which the cue was on. If animals touched the cue during this period, a reward three times the size of the regular reward was dispensed (840 ms). This led to rapid acquisition. Following this, animals were exposed daily to a 30-min session of must touch training. This schedule followed the same procedure as the initial touch training, but free reinforcers were terminated and animals were required to nose poke the image in order to obtain a regular reward (7-uL, 280 ms).

#### Deterministic pairwise discrimination training

Animals were exposed to 10 days of pairwise discrimination training, during which animals were presented with two highly discriminable image cues (“marbles” and “fan”). One image was always rewarded and the other one was not. Within each session, animals completed either 250 trials or spent a maximum of two hours in the operant chamber (typically these mice completed ∼200 trials/day).

#### Two-armed bandit task

Animals were trained to perform a two-arm visual bandit task in the touchscreen operant chamber. Each trial, animals were presented with a repeating set of two different images on the left and right side of screen, counterbalancing left and right across the session. Nose poke to one of the displayed images on the touchscreen was required to register a response. Nose poke on one image triggered a reward 80% of the time (high payoff image), whereas the other image was only reinforced 20% of the time (low payoff image). Following the reward collection, which was registered as entry and exit of the feeder hole, the magazine would illuminate again and the mouse must re-enter and exit the feeder hole to initiate the next image trial. If the previous trial was unrewarded, a 3-second time-out was triggered, during which no action could be taken. Following the timeout, the magazine would illuminate and the mouse must enter and exit the feeder hole to initiate the next image trial. The ABET II system recorded trial to trial image chosen history, reward history, grid position of the images with time-stamp. Within each day, animals completed either 250 trials or spent a maximum of two hours in the operant chamber. Animals were given 14 days to learn about the probabilistic reward schedule of one image pair, before moving onto the next image pair. A total of six image pairs were trained, but two image pairs were eliminated from analyses due to very high initial preference (>70%) for one novel image over another, indicating that (to the mice) these images appeared unexpectedly similar to previously experienced images with learned reward values.

#### RNA quantification

At the end of training, animals were sacrificed after the second day of learning a new image pair (around 400-500 trials of experience per mouse), when we expected to see the biggest difference in learning performance and strength of lateralization. Animal brains were extracted and targeted brain regions were dissected. We extracted RNA from targeted brain areas and assessed gene expression for the *fos* genes in the nucleus accumbens (NAc), dorsal medial striatum (DMS), amygdala (AMY), and hippocampus (HPC), using quantitative Real Time PCR system (BioRad, USA). Fos expression normalized to the housekeeper gene glyceraldehyde 3-phosphate dehydrogenase (*gapdh*) was calculated using the comparative delta Ct method.

### Data analysis

#### Generalized Linear Models (GLMs)

In order to determine whether sex and number of trials (bins) predicts the accuracy of the task, strength of lateralization, reaction time, mutual information (MI), or angle between probability vectors, we fit a series of generalized linear models of the following form:

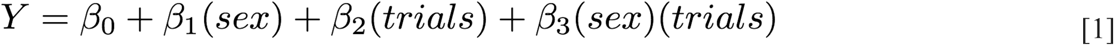

Where Y is the dependent variable (accuracy, laterality, reaction time, MI, or angle). In this model, β1 described the main effect of sex and β2 described the main effect of number of trials (bins). β3 captures any interaction effect between sex and number of trials (bins).

To determine whether c-fos expression in NAc, DMS, AMY, HPC, PFC, and sex predicted the weights of Principal Component (PC) 2, we fit the following generalized linear model.

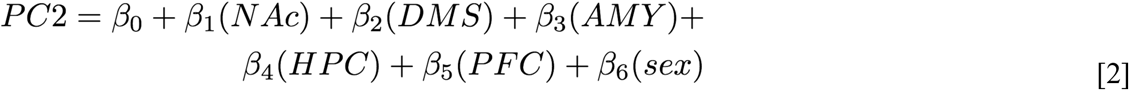

In this model, β1-β5 captures the predictive effect of gene expression in five regions on the use of PC2 strategy. β6 described the effect of sex on the weights of PC2.

#### Degree of lateralization

As a measure of the strength of side bias, we used the absolute percentage of laterality [39], calculated for each mouse according to the following formula:

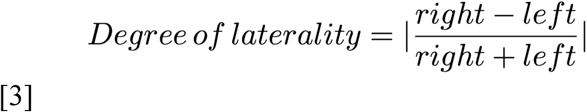

#### Generalized Logistic Regression Model

Mice could base their decisions on reward history in the spatial or image domains or on choice history in the spatial or image domains. To determine how these four aspects of previous experience affected choice and how these effects changed over time, we estimated the effect of the last trials’ reward outcome (O), image choice (I), and chosen side (S) using logistic regression. If image (image 1) was on the left side of the screen, we could predict the probability of choosing that image as a linear combination of the following four terms:

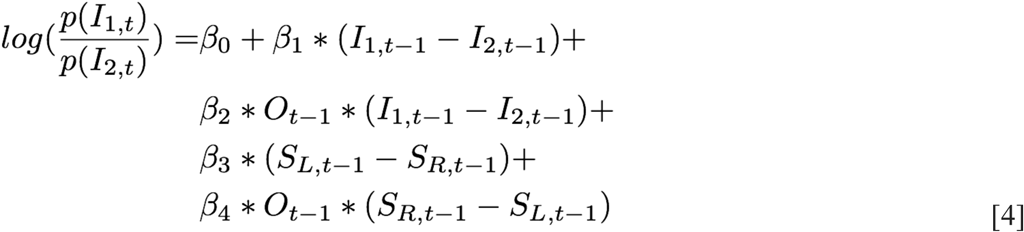

Where each term (O, I, and S) is a logical, indicating whether or not that event occurred on the last trial. As a result, the term (I_{1, t-1} - I_{2, t-1}) is 1 if image 1 was chosen on the last trial, but −1 otherwise. The term β1 thus captures the tendency to either repeat the previous image (when positive) or choose the other image (when negative). The term β2 O_{t-1} accounts for any additional effect of the previous image on choice, when that previous choice was rewarded.. If image 1 was on the left side, (S_{L, t-1} denotes the probability of repeating the left side where image 1 appeared. However, because image 1 could be either on the left or the right side of the screen (which allowed us to dissociably estimate the probability of choosing it based on side bias or image bias), we expanded the (S_{L, t-1} - S_{R, t-1}) term to account for the current position of image 1 as follows:

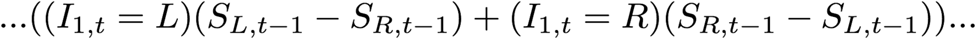

Meaning that the current position of image 1 determined the sign of the side bias term. This model was fit individually to each bin of 150 trials, within each animal and image pair, via cross-entropy minimization with a regularization term (L2/ridge regression).

#### Principal component analysis

In order to determine how decision-making strategies differed across animals and bins, we looked for the major dimensions of interindividual variability in decision-making strategies. To do this, we took advantage of the fact that the coefficients of the generalized linear model provided a simplified description of how decision-making depended on image, side, and outcome for each subject in each bin. Because the generalized logistic regression model estimated 4 terms per image pair and there were 23 independent bins per image pair, this meant that each animals’ behavior for a given image pair could be described as a 4*23 by 1 dimensional vector. We then used principal component analysis to identify the linear combinations of model parameters that explained the most variance across subjects and repetitions of image pairs (across 32 (animals) × 4 (repetitions) = 128 total strategy vectors). The first two principal components, which explained the majority of the variance (59%), are illustrated in Figure 2e.

#### Conditional mutual information and model-free analyses

To account for idiosyncratic strategies, which could vary across animals or image pairs, we used a model-free approach to quantify the extent to which behavior was structured without making strong assumptions about what form this structure might take. We quantified the extent to which choice history was informative about current choices as the conditional mutual information between the current choice (C) and the last choice (C_t-1), conditioned on the reward outcome of the last trial (R):

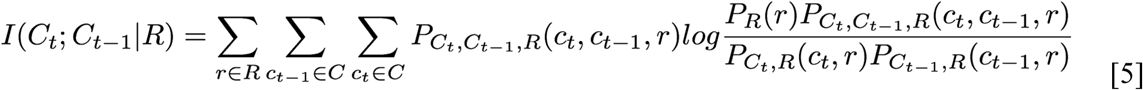

Where the set of choice options (C) represented the unique combinations of each of the 2 images and 2 sides (4 combinations). To account for observed differences in overall probability of reward for male and female animals, the mutual information was calculated independently for trials following reward delivery and omission, and then summed across these two conditions.

We used a similar approach to provide a model-free description of the animals’ choice patterns. Briefly, instead of finding the set of beta weights that best described reliance on various history-dependent strategies over time, we directly calculated the joint probability of each possibility combination of last choice (image and side), last outcome (reward and unrewarded), and current choice (image and side). This means that we represented the animals’ history-dependent choice pattern for each image pair as an 32-dimensional vector (4 (last choice) × 2 (last outcome) × 4 (current choice) = 32) of joint probabilities. Via a geometric interpretation of a multinomial distribution, we considered the animal’s pattern of behavior within any bin of trials as a point on the 32-1 dimensional simplex formed by length-1 vectors. This geometric approach allowed us to map strategies over time or across bins as a diffusion process across this simplex, where the angle between two vectors (between animals/between bins/between repetitions) is proportional to step between them on a strategy simplex. The bigger the step between two vectors, the more variable the behavior pattern is.

#### Mediation Analysis

First we used a direct model and regressed c-fos expression of either NAc or PFC on weights of PC2. When assessing a mediation effect, three regression models are examined:

Model 1 (direct):

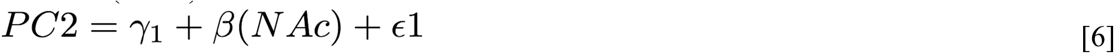

Model 2 (mediation):

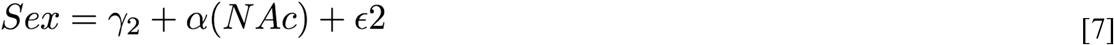

Model 3 (indirect)

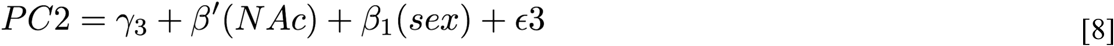

In these models, γ1, γ2, and γ3 represent the intercepts for each model, while ε1, ε2, and ε3 represent the error term. β denotes the relationship between dependent variable (PC2 weights) and independent variable (NAc c-fos expression) in the first model, and β’denotes the same relationship in the third model. α represents the relationship between independent variable (NAc c-fos expression) and mediator (sex) in the second model. The mediation effect is calculated using the product of coefficients (αβ1). The Sobel test is used to determine whether the mediation effect is statistically significant [46].

#### Reinforcement Learning Model

We considered a basic reinforcement learning model, following the Rescorla Wagner rule. In this model, subjects first learn the expected value of each image based on the history of its previous outcome value Q and use these Q values to decide what to do next. The expected value of arm *k* on the *t* th trial, *Q*_*t*_ ^*k*^, is updated based on the reward outcome of each trial:

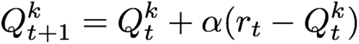

In each trial, r_t_ – Q _t_ ^k^ captures the reward prediction error (RPE), which is the diffe-rence between expected outcome and the actual outcome. The parameter *a* is the learning rate, which determines the rate of updating RPE. Action selection was performed based on a Softmax probability distribution:

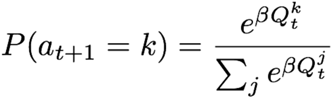

where inverse temperature *b* determines the level of random exploration.

## Acknowledgements

The authors would like to thank Sarah Heilbronner and Vincent Costa for helpful comments on the manuscript. This work was supported by NIMH T32 training grant MH115886, startup funds from the University of Minnesota (NMG), and a NARSAD Young Investigator Grant (RBE).

## Notes

#### Summary of Updates

Abstract and introduction updated

